# Closed-loop Control of Functional Electrical Stimulation Using a Selectively Recording and Bidirectional Nerve Cuff Interface

**DOI:** 10.1101/2023.06.22.546126

**Authors:** Y. E. Hwang, L. Long, J. Sales Filho, R. Genov, J. Zariffa

## Abstract

**Background:** Discriminating recorded afferent neural information can provide sensory feedback for closed-loop control of functional electrical stimulation, which restores movement to paralyzed limbs. Previous work achieved state-of-the-art off-line classification of electrical activity in different neural pathways recorded by a multi-contact nerve cuff electrode, by applying deep learning to spatiotemporal neural patterns.

**Objective:** To incorporate this approach into closed-loop stimulation.

**Methods:** Acute *in vivo* experiments were conducted on 11 Long Evans rats to demonstrate closed-loop stimulation. A 64-channel (8 × 8) nerve cuff electrode was implanted on each rat’s sciatic nerve for recording and stimulation. A convolutional neural network (CNN) was trained with spatiotemporal signal recordings associated with 3 different states of the hindpaw (dorsiflexion, plantarflexion, and pricking of the heel). After training, firing rates were reconstructed from the classifier outputs for each of the three target classes. A rule-based closed-loop controller was implemented to produce ankle movement trajectories using neural stimulation, based on the classified nerve recordings. Closed-loop stimulation was initiated by the detection of a heel prick, and induced dorsiflexion. The detection of dorsiflexion triggered stimulation to induce plantarflexion, and vice versa. A single trial began with a heel prick and ended when an incorrect state transition occurred or when a second heel prick was detected.

**Results:** Closed-loop stimulation was successfully demonstrated in 6 subjects. Number of successful trials per subject ranged from 1-17 and number of correct state transitions per trial ranged from 3-53.

**Conclusion:** This work demonstrates that a CNN applied to multi-contact nerve cuff recordings can be used for closed-loop control of functional electrical stimulation.

## 1 Introduction

Spinal cord injury can disrupt communication between the central nervous system and the periphery, resulting in impairment of limb movements previously enabled by nuanced supraspinal motor control signals. The restoration of motor function in paralyzed limbs can be achieved using functional electrical stimulation (FES), which triggers desired muscle contractions through patterned electrical stimulation applied to motor axons. Replicating the sophistication of motor commands generated by the CNS using pre-programmed open-loop FES is challenging. This form of FES delivery typically yields limb movement that lacks nuance and the ability to react to unexpected perturbations or errors, and does not provide an efficient delivery of stimulation that avoids excessive muscle fatigue [1]. These shortcomings can be addressed by closed-loop FES, which uses sensor feedback to guide stimulation delivery parameters such as stimulation timing, target location, frequency, and intensity. Afferent neural information is an appealing avenue for extracting sensory feedback for closed-loop FES, because it avoids the need for external sensors and directly encodes proprioceptive and sensory information used for motor control [2].

Peripheral nerve interfaces can be used to record afferent signals and extract proprioceptive and sensory information through data processing. Closed-loop FES using afferent nerve signals as feedback has been demonstrated in both animal and human trials. For example, Bruns et al. and Holinski et al. both demonstrated FES for cat leg motion control using activity recorded in the dorsal root ganglion (DRG) and decoded by linear regression and multivariate linear equations, respectively [3] [4]. Song et al. [5] demonstrated FES for rabbit ankle motion control using activity recorded in the sciatic nerve and decoded using fast independent component analysis combined with a dynamically driven recurrent neural network. Inmann et al. [6] demonstrated hand grasp control in humans using activity recorded from the median nerve decoded by rectification and bin-integration, signal filtering, and delay.

Closed-loop FES in clinical applications should use highly selective recording and stimulation techniques and be implemented on devices that are as minimally invasive as possible. Interfacing with multiple individual nerve branches instead of their originating nerve trunk allows for superior selectivity but increases invasiveness due to multiple surgical implants. Similarly, using intraneural or regenerative electrodes allows for superior selectivity compared to extraneural electrodes at the cost of invasiveness [7], [8]. Apart from Song et al., each aforementioned study chooses selectivity over minimal invasiveness for one or more of these trade-offs, so they are not yet ideal for translation to chronic practical use.

Koh et al. demonstrated that a convolutional neural network (CNN) could achieve state-of-the-art accuracy for classifying nerve recordings obtained by multi-contact extraneural nerve cuff electrodes, a method named Extraneural Spatiotemporal Compound Action Potentials Extraction Network (ESCAPE-NET) [9]. Each electrode contained multiple neural interfacing contacts around the circumference of the sciatic nerve organized as uniformly separated rings along the length of the nerve to capture spatial and velocity variability between different neural pathways. Individual naturally-evoked compound action potentials (nCAPs) were detected and represented as an image referred to as the spatiotemporal signature, which the CNN was trained to classify. An advantage of classifying individual nCAPs is to provide high temporal resolution that could potentially handle the scenario of multiple simultaneously active neural pathways, because individual nCAPs are less likely to overlap in time than when averaging over longer time windows.

The results reported by Koh et al. were based on an off-line analysis of the recorded signals [9], without the presence of stimulation. The objective of this study is to demonstrate that the ESCAPE-NET classification method can be used to provide closed-loop feedback for modulating nerve stimulation. We implement a closed-loop FES system based on a bidirectional, selective peripheral neural interface using a single multi-contact nerve cuff. The novel contribution of this work is to demonstrate that extraneural classification of individual nCAPs is feasible in a closed-loop FES scenario.

## 2 Methods

To demonstrate ESCAPE-NET’s applicability for closed-loop FES, a closed-loop system was designed to produce a movement pattern in response to the selectively monitored afferent activity of several neural pathways. A simple lower limb alternating movement task was selected to validate the system while taking advantage of a well-established experimental model for selective peripheral nerve interfacing, the rat sciatic nerve.

### 2.1 Surgical approach

Acute experiments were conducted on 11 Long-Evans rats (Envigo, Indianapolis, USA). The procedures were approved by the Animal Care Committee of the University Health Network (#6477). Retired male breeders were used due to size requirements for the electrode implantation. Anesthesia was induced through inhalation using isoflurane. The procedure began with an oblique incision along the posterior and dorsal aspect of the hip. Connective tissue was cleared around the incision area to reveal the biceps femoris and gluteus maximus muscles [10], [11], which when separated provide access to the sciatic nerve. The nerve was then gently cleaned of connective tissue using dull instruments, being careful not to damage or sever any nerve matter. A custom-made planar polyimide electrode was folded over the sciatic nerve proximally to the bifurcation point into the tibial, peroneal and sural nerves, avoiding excessive stretching or scratching of the nerves. The electrode was then fixed in place by stitching through its suture holes. An Ambu Neuroline Monopolar Electromyography needle electrode was placed subcutaneously into the rat’s back as a ground.

The electrode used was an extraneural nerve cuff with 64 contacts arranged in 8 rows of 8 contacts. Figure 1 illustrates the electrode panel design and contact layout. The electrode folds over the nerve along the horizontal axis, bending along the space between the fourth and fifth row of contacts. The electrode is 20 mm long along the horizontal axis and 8 mm long along the vertical axis. Contacts are 0.75 mm long along the horizontal axis and 0.25 mm long along the vertical axis. The electrodes were manufactured using commercial fabrication methods on a 2-layer polyimide (25 micron thickness) flex printed circuit board with a chemical gold ENIG finish (copper thickness 25 μm, nickel thickness 4 μm, gold thickness 0.05 μm). The reference contact was implemented as a 1 mm x 1 mm contact on the outside of the cuff.

**Figure 1:**
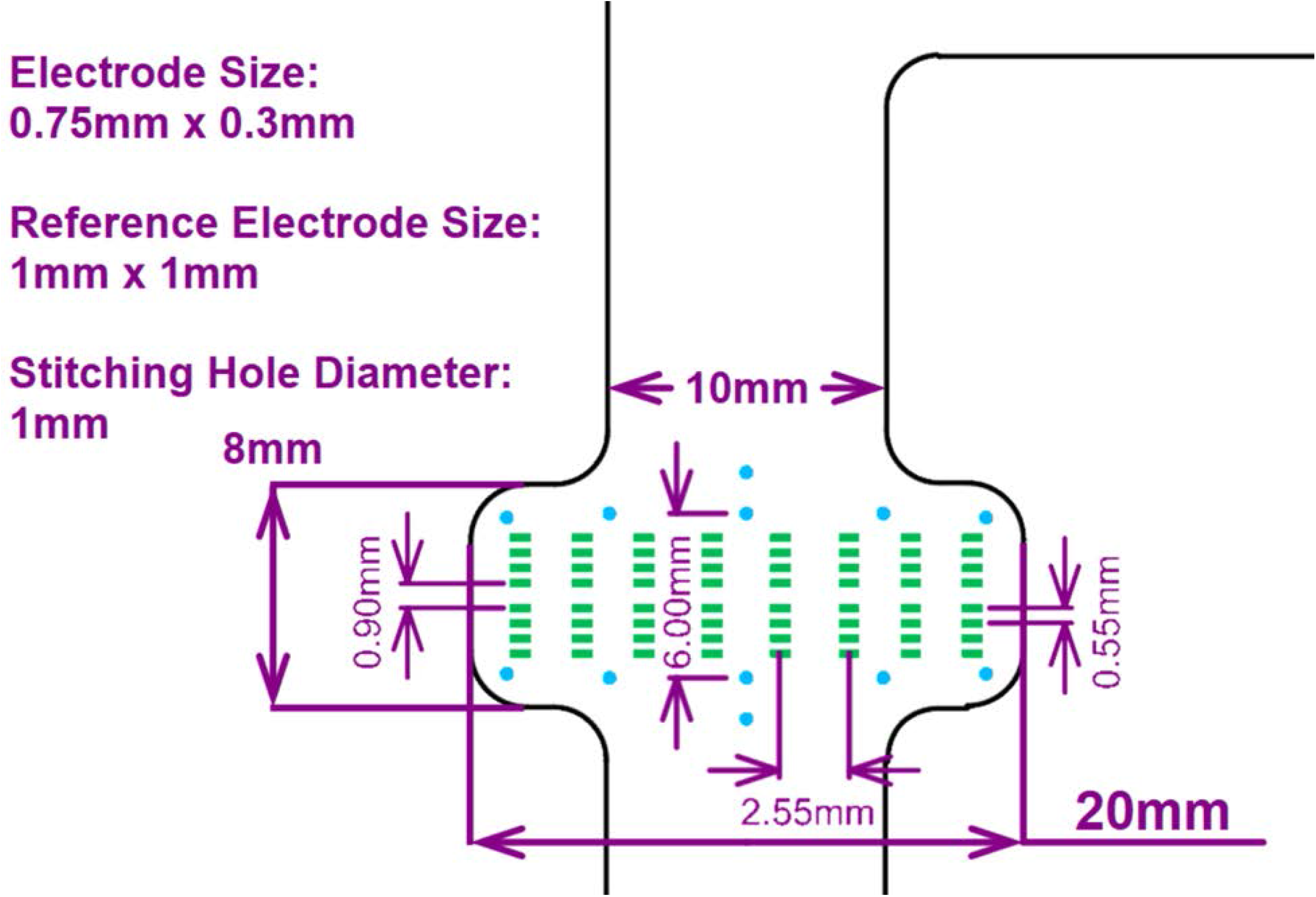
Schematic of the electrode’s contacts layout. The electrode is folded over the nerve along the horizontal axis. Each column of contacts is pressed against the circumference of the nerve, and the 8 columns are spaced along the length of the nerve. Suturing through the stitching holes (6 blue holes at the top and bottom of the panel) secures the electrode in position. A reference contact is situated on the backside of the depicted surface.

### 2.2 Closed-loop stimulation

To detect and classify afferent nCAPs using ESCAPE-NET, the closed-loop system sequentially performed the following operations: data acquisition, signal pre-processing (including stimulation artefact removal), nCAP detection, spatiotemporal signature conversion, and ESCAPE-NET classification of spatiotemporal signatures. To use the results of online nCAP classification to modulate stimulation parameters, the system performed the following additional components: neural pathway firing rate monitoring, and a rule-based controller for reactive stimulation. These steps are individually described in the following sections.

### 2.3 Detecting and classifying nCAPs using ESCAPE-NET

#### 2.3.1 Data acquisition, signal pre-processing, and nCAP detection

Signals were recorded by the nerve electrodes with a sampling rate of 30kHz (CerePlex Direct, Blackrock Neurotech, Salt Lake City, UT, USA). The obtained raw signals underwent signal processing to detect and represent nCAPs in a consistent format for classification. An nCAP is the summation of action potentials firing synchronously from a group of neurons with similar functional purposes, evoked by naturally occurring phenomena such as changes in limb position or tactile input (as opposed to a CAP evoked by direct electrical stimulation of the nerve). An nCAP corresponds to a detectable “spike” in the nerve cuff electroneurogram (ENG) signal. In this study, an nCAP was classified as tibial branch activity (induced by dorsiflexion of the ankle [12]), peroneal branch activity (induced by plantarflexion of the ankle [12]), or sural branch activity (induced by applying a cutaneous stimulus to the heel using a Von Frey monofilament [300 g] [13]).

Dorsiflexion and plantarflexion of the ankle were administered both manually and through electrical stimulation of the sciatic nerve. Stimulation was delivered (CereStim R96, Blackrock Neurotech, Salt Lake City, UT, USA) to the sciatic nerve from each contact of the nerve cuff, and contact selection was performed manually based on the selectivity and amplitude of movement for both dorsiflexion and plantarflexion. Based on the subject, stimulation sometimes needed to be delivered to multiple contacts simultaneously to induce clear movement. Contact selection was an iterative process as initially effective contacts could lose effectiveness throughout the experiment. Stimulation was delivered in bursts of 30 biphasic pulses, with amplitude of 215 μA, frequency of 30Hz, phase width of 100 μs, and interphase delay of 100 μs.

To remove any artefacts caused by electrical stimulation of the nerve, the recorded signal was zeroed for a portion of time extending from immediately before (0 – 1 ms) to after (10 – 11 ms) a stimulation pulse. Tripolar referencing was applied to the acquired raw signal, using the average of the contacts in the two outer rings as the reference. Bandpass filtering was then applied using a 4th order Butterworth filter with a 1–3 kHz passband. nCAPs were detected by using the delay-and-add method [14]–[18] (applied to the averaged signal of each ring of contacts, centered around either the 4^th^ or 5^th^ ring of contacts, determined empirically) to obtain a superimposed nCAP. The delay-and-add method uses conduction velocity and contact spacing to temporally align nCAPs recorded by longitudinal contacts to obtain a larger superimposed nCAP that is easier to detect. An nCAP was detected anytime the superimposed nCAP signal X crossed a threshold determined by equation (1). To eliminate artefactual data, any nCAP that crossed a threshold of 20 μV was discarded. In this manner, afferent nCAPs were extracted from the intervals between stimulation pulses.

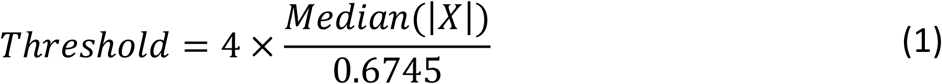

#### 2.3.2 Spatiotemporal signature

After each detected nCAP was tripole referenced, it was used to create a spatiotemporal signature representation, using 49 time samples before and 50 time samples after the superimposed nCAP’s peak, corresponding to 3.33 ms segment centered around the detected spike [9]. Note that the delay-and-add operation was used only to detect nCAPs, but the spatiotemporal signatures were extracted from the 64-channel signals. The spatiotemporal signature is an M × T matrix, with M contacts on the electrode and T time samples (here 64 and 100, respectively). The grid of contacts on the electrode was numbered to determine the ordering in the M dimension, as detailed in the “ESCAPE-NET Architecture” section below. Spatiotemporal signatures were used as inputs to the CNN.

#### 2.3.3 ESCAPE-NET training

ESCAPE-NET was trained using isolated trials of sciatic nerve afferent activity induced by applying dorsiflexion (tibial branch activity) and plantarflexion (peroneal branch activity) to the rat’s ankle [12] and applying a cutaneous stimulus to the heel (sural branch activity) using a Von Frey monofilament (300 g) [13]. A training dataset unique to each specific subject was obtained. Dorsiflexion and plantarflexion were induced both manually and through electrical stimulation; pricking of the heel was induced manually. The inclusion of electrically stimulated movement in the training set was motivated by the fact that the nCAPs detected in the closed-loop demonstration are electrically stimulated. Offline training was performed remotely on a desktop computer equipped with a GeForce RTX 3090 GPU during the *in vivo* experiment.

100 trials of each of the 3 types of manual stimuli were initially performed. As the process of data transfer, signal pre-processing, conversion to training dataset, and NN training typically took over an hour, additional training was continually performed on new smaller datasets after the initial training stage. Retraining was performed to adapt the NN to any minor shifts of the electrode’s positioning or changes in the recording site’s interface with the tissue. Performing additional training on a new smaller dataset was much less time-intensive, typically taking a few minutes. Across all subjects, up to 6 iterations of retraining were performed. The additional training dataset was comprised of activity induced by electrically stimulated dorsiflexion and plantarflexion, and manual pricking of the heel. The number of trials for each type of activity during retraining was empirically adjusted to ensure similar numbers of detected nCAPs across all three labels, typically aiming to achieve 100-200 detected nCAPs.

#### 2.3.4 ESCAPE-NET architecture

A modified version of ESCAPE-NET [9] was used to classify detected nCAPs as illustrated in Figure *2*. The CNN receives two inputs, corresponding to two different representations of the spatiotemporal signature. The two representations differ in the ordering of the contacts. The “spatial emphasis” representation orders contacts by ring (i.e. every 8 rows of the image belong to a ring of contacts), which ensures that rows of pixels are locally related based on contact placement around the circumference of the nerve; the “temporal emphasis” representation orders contacts by length (i.e. every 8 rows of the image belong to a line of contacts running along the nerve), which ensures that rows of pixels are locally related based on contact placement along the length of the nerve. The former exploits the spatial variation of different neural pathways, and the latter exploits the velocity variation of different neural pathways.

For each of the two inputs of the network, there are 3 convolutional layers in sequence that have 32 filters and use the ReLU activation function. The first convolutional layer has 8×8 filters, the second has 4×4 filters, and the third has 2×2 filters. The convolutional layers are separated by 2×2 max pooling layers for data downsampling. To initialize the weights of the first and second convolutional layers, they were temporarily connected to dense layers and trained for 25 epochs. The pair of convolutional layer stacks are concatenated after the third convolutional layer into a dense layer that has 64 neurons and uses the ReLU activation function. Dropout at a rate of 0.5 is used on this dense layer. The final layer is a dense layer responsible for classification into the 3 possible classes.

This version of ESCAPE-NET uses 2,912,963 weights and 150,013,714 floating point operations required for interference on a single input. Possible optimizations of this neural network architecture have recently been investigated to make it appropriate for use in implanted systems [19], [20] however for these *in vivo* experiments the architecture was kept fixed.

**Figure 2:**
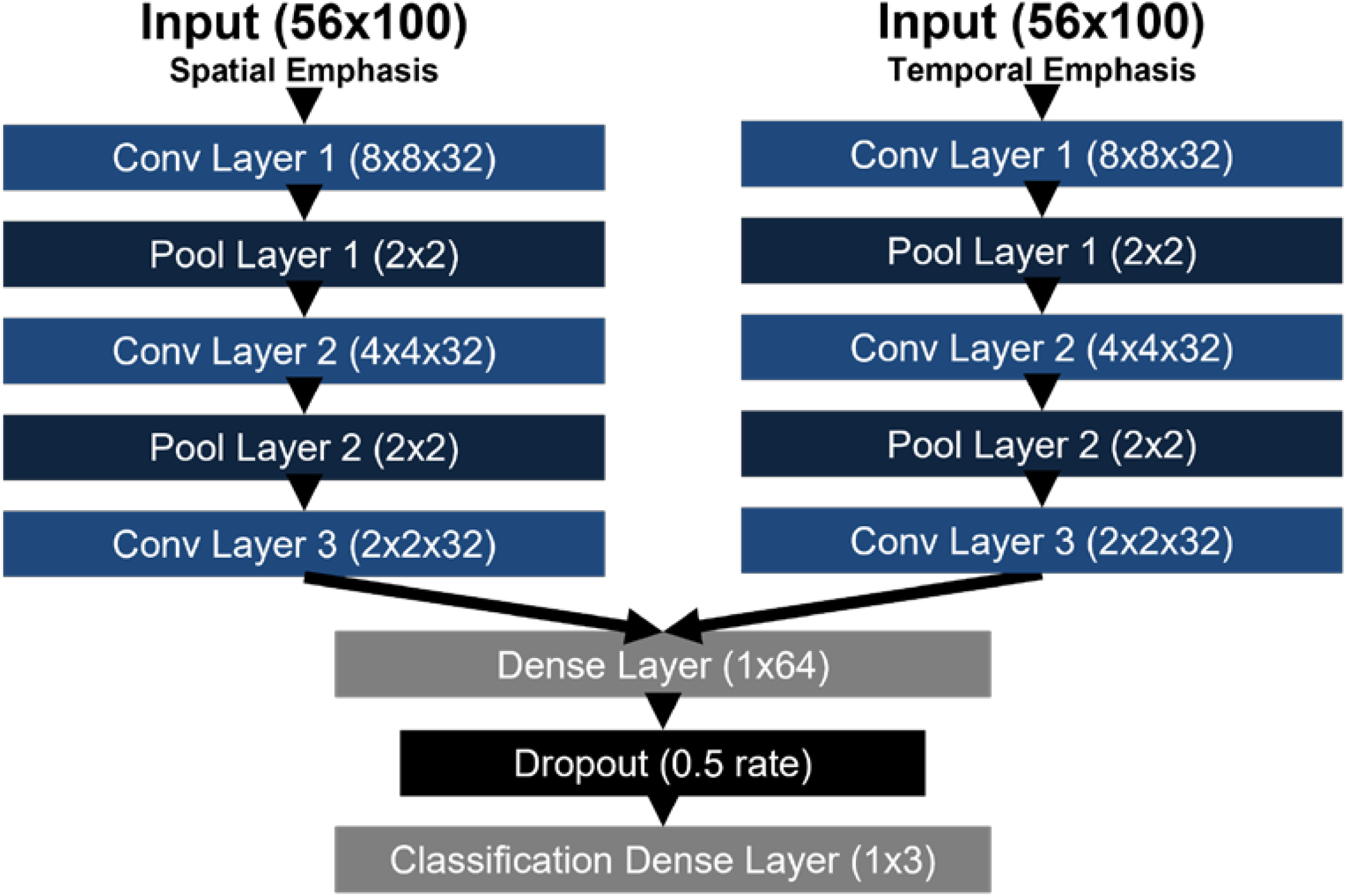
Illustration of a modified version of ESCAPE-NET, the CNN used for classifying spatiotemporal signatures.

### 2.4 Using online nCAP classifications to modulate stimulation parameters

All components were implemented online in MATLAB, using the Signal Processing toolbox for pre-processing, the Deep Learning toolbox for Keras NN classification, and Blackrock’s MATLAB interfaces cbMEX and StimMex to receive recorded data and control stimulation, respectively. Figure *3* illustrates the devices used for the closed-loop stimulation demonstration and the interactions between them. The experimental setup (online setting) included a laptop computer equipped with MATLAB which implemented a rule-based controller that directly interacted with the data acquisition system and stimulation device. The data acquisition system and stimulation device were both connected to the custom electrode, which was surgically implanted in the subject. The laptop computer had remote access to a desktop computer equipped with a GeForce RTX 3090 GPU for neural network training, allowing for dataset acquisition, neural network training, and closed-loop stimulation to be performed on the same subject within one experiment session. The latency of the closed-loop system was empirically observed to be approximately 3 seconds, which is discussed in more detail in the Discussion section.

**Figure 3:**
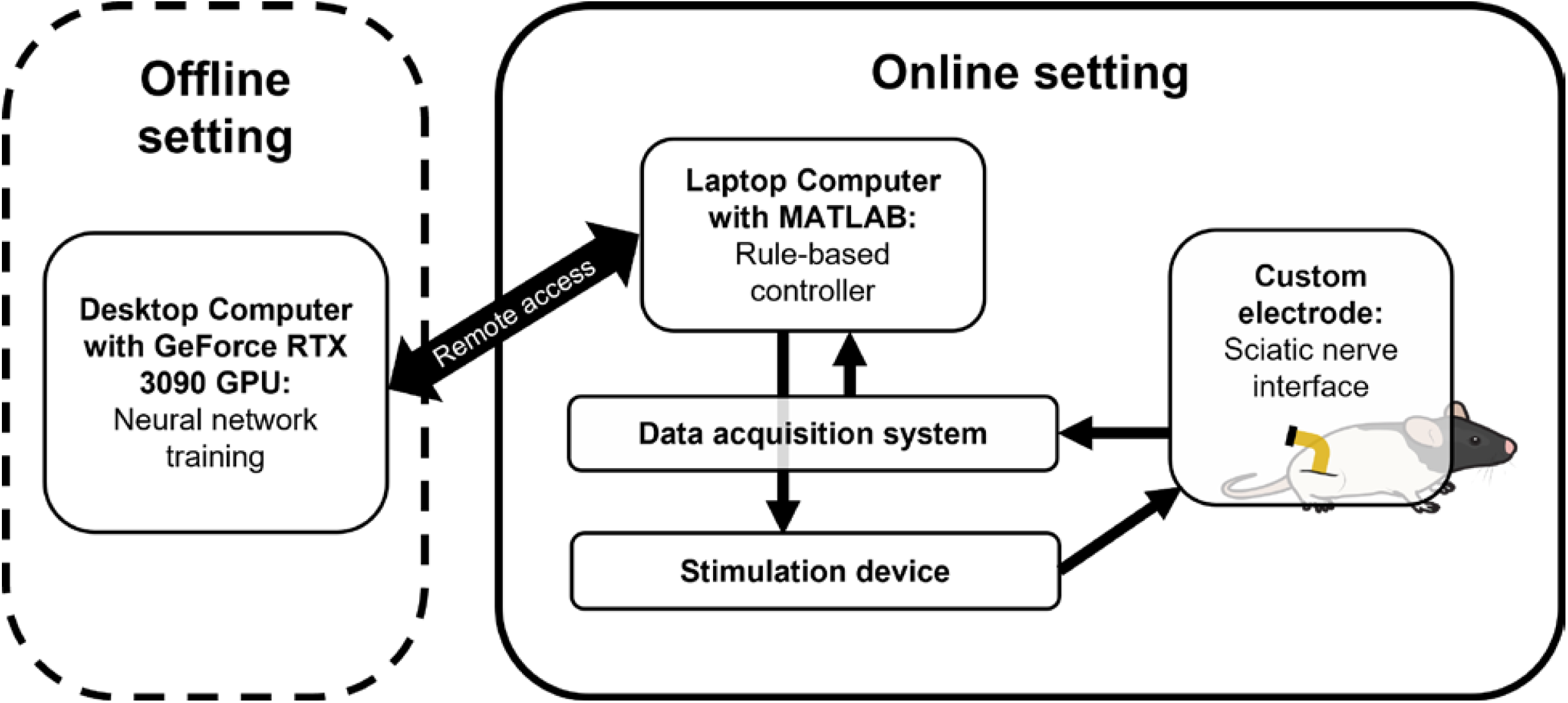
Diagram illustrating the experimental setup for in vivo closed-loop stimulation demonstration (rat image from [21]).

#### 2.4.1 Neural pathway firing rate monitoring

The peroneal, tibial and sural branches’ firing rates were computed by detecting and classifying nCAPs using ESCAPE-NET within approximately 3-second segments of continuously acquired signal, with 50 samples (1.67 ms) of overlap in between windows to capture any nCAPs that are cut off between windows. For clarification, windows were used so that finite segments of signal could be analyzed in a continuous fashion and firing rates computed – our method classifies individual nCAPs, rather than windows of activity.

#### 2.4.2 Rule-based controller design for closed-loop stimulation

Controller design for closed-loop FES is a complex topic, for which a number of sophisticated strategies have been proposed [22]–[25]. In order to de-couple this complexity from the objective of this experiment, which was to demonstrate classification of individual nCAPs in a closed-loop FES scenario, we designed a simplified control paradigm based on finite state machines. The task was designed to minimize controller complexity while requiring two key elements: selective recording from multiple neural pathways, and interleaved recording and stimulation from a single electrode.

Active branches were used to transition between different states of stimulation. Figure *4* illustrates finite state machines (FSM) describing the possible states of stimulation that were administered, and which active branch triggered a transition between states. We refer to the FSM depicted in Figure *4*a as FSM A. Figure *4*b depicts a slightly more sophisticated version of the FSM, which we will refer to as FSM B. Initial focus was placed on demonstrating FSM A, and once it was demonstrated consistently across multiple subjects, both FSMs A and B were attempted. The FSMs were simplified movement tasks selected as an experimental model to reflect a clinically meaningful scenario (monitoring of both joint angles and external stimuli for closed-loop control of FES) while requiring selective neural recording.

**Figure 4:**
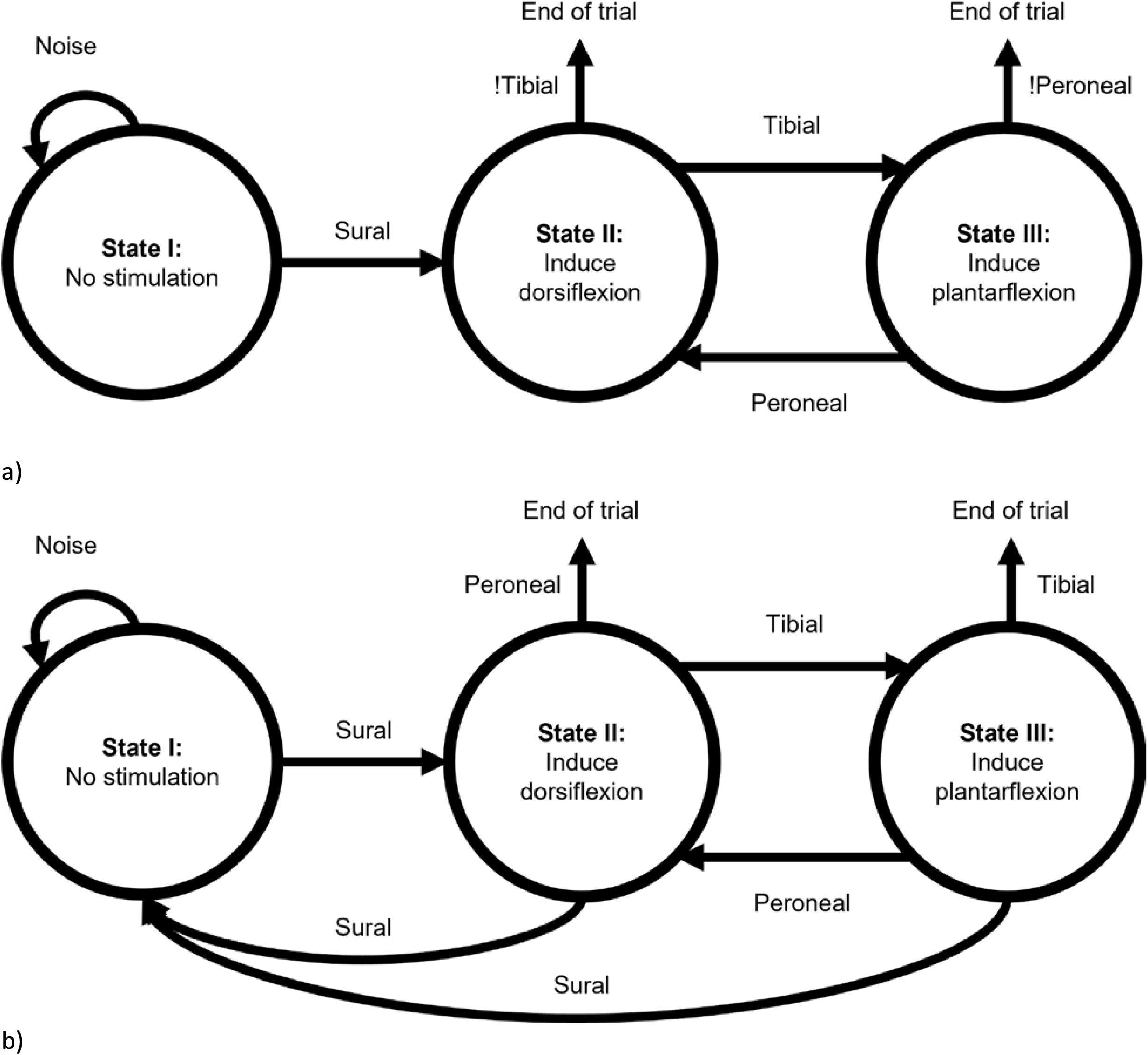
Diagram of the FSMs used in the rule-based controllers for closed-loop stimulation. a) Diagram depicting FSM A, which ends when a window of activity is incorrectly classified. B) Diagram depicting FSM B, which ends when a heel prick is detected, or a window of activity is incorrectly classified.

The desired movement pattern was initiated with a prick of the heel, then oscillating ankle motions was induced via nerve stimulation. In FSM A, oscillation continued until a mistake in monitoring occurred, determined by either tibial or peroneal activity being detected in two consecutive windows. For the FSM A experiments, the highest firing rate within a window was used to identify the active branch (simultaneous activations were not considered). The experiment began in state I, with the foot at rest in a neutral position. No stimulation was administered in state I. If only noise was detected (i.e. the total number of nCAPs was below the noise threshold), state I continued to be held. When a heel prick (sural branch activity) was detected, the state transitioned to state II, which selectively stimulates the nerve to induce dorsiflexion. When full dorsiflexion (tibial branch activity) was detected, the state transitioned to state III, which selectively stimulated the nerve to induce plantarflexion. When full plantarflexion (peroneal branch activity) was detected, the state transitioned back to state II.

In FSM B, oscillation continued until a second prick of the heel was detected. This required detecting activity in multiple branches simultaneously (both proprioceptive and cutaneous afferents). To detect multiple simultaneously active branches, a firing rate threshold per window was defined for each nerve branch. This was done by analyzing successful trials of FSM A, and choosing a threshold value for each of the three types of neural activity that would yield correct activity detection.

## 3 Results

### 3.1 Selective recording during closed-loop FES

Closed-loop stimulation was successfully demonstrated in six subjects. Table 1 reports the number of successful trials achieved for all subjects for which closed-loop stimulation was attempted. A successful trial is defined as one where all 3 neural pathways were correctly identified as active at least for one window each. In other words, each possible transition between states of the FSM should be performed at least once. For each successful trial, the number of transitions between states was recorded to quantify the robustness of a single trial. The mean and standard deviation for number of state transitions is also reported.

**Table 1:** Summary of trials achieved for subjects with successfully demonstrated closed-loop stimulation. N/A indicates that a protocol was not attempted.

| Subject | FSM A:<br>Number of trials | FSM A:<br>Number of state transitions |  | FSM B:<br>Number of trials | FSM B:<br>Number of state transitions |  |
| --- | --- | --- | --- | --- | --- | --- |
| | | Per trial | Mean ± standard deviation | | Per trial | Mean ± standard deviation |
| 1 | 0 | N/A |  | 0 | N/A |  |
| 2 | 0 | N/A |  | 0 | N/A |  |
| 3 | 0 | N/A |  | 0 | N/A |  |
| 4 (A) | 5 | 10, 10, 10, 8, 9 | $9.4 \pm 0.8$ | N/A | N/A | |
| 5 (B) | 17 | 8, 4, 4, 3, 4, 4, 12, 8, 4, 10, 14, 23, 3, 10, 3, 6, 11 | $7.7 \pm 5.2$ | N/A | N/A | |
| 6 (C) | 12 | 4, 6, 13, 25, 21, 25, 37, 37, 31, 53, 43, 41 | $28 \pm 14.6$ | N/A | N/A | |
| 7 (D) | 7 | 16, 10, 5, 15, 7, 3, 4 | $8.6 \pm 4.9$ | 2 | 4, 5 | $4.5 \pm 0.5$ |
| 8 | 0 | N/A |  | 0 | N/A |  |
| 9 (E) | 1 | 9 | $9 \pm 0$ | 0 | N/A | |
| 10 | 0 | N/A |  | 0 | N/A |  |
| 11 (F) | 6 | 4, 40, 4, 4, 4, 4 | $10 \pm 13.4$ | 7 | 4, 4, 5, 4, 5, 5, 7 | $4.9 \pm 1.0$ |

Retraining of the network using new smaller datasets occurred between certain trials within a single subject, so the weights of a network used between trials of a subject are not necessarily the same. Also, contact configurations for stimulation delivery often changed over the course of an experiment. Trials of FSM B were not attempted on subjects A-C, and were attempted on subjects D, E and F; trials of FSM B were not achieved for subject E.

Figure *5* shows an example of the firing rate trajectories of the three monitored nerve branches within a trial compared to the expected active branch based on the position of the rat’s foot and presence of cutaneous stimulus to the heel. In Figure *5*a, the highest firing rate per window corresponds to the expected active branch. As dictated by FSM A, detection of sural branch activity begins stimulation delivery, inducing oscillating movements between dorsiflexion and plantarflexion. In Figure *5*b, sural branch activity is detected using threshold crossing, whereas tibial and peroneal branch activity remains being detected based on the highest firing rate; by doing so, sural branch activity is monitored separately, and can be used to override stimulation decision-making. As dictated by FSM B, detection of sural branch activity begins stimulation delivery as in FSM A, and a second detection of sural branch activity halts stimulation delivery. One undetected instance of expected sural branch activity occurred at approximately the 36 s mark. Although sural branch activity detection halted stimulation delivery, tibial branch activity was still successfully detected within the same window. Note that the ground truth and detected activity events in Figure 5 are not completely aligned because the window length used to estimate firing rates was longer than the sensory events. Nonetheless, the variations in the relative firing rates demonstrate the responsiveness to these sensory events.

**Figure 5:**
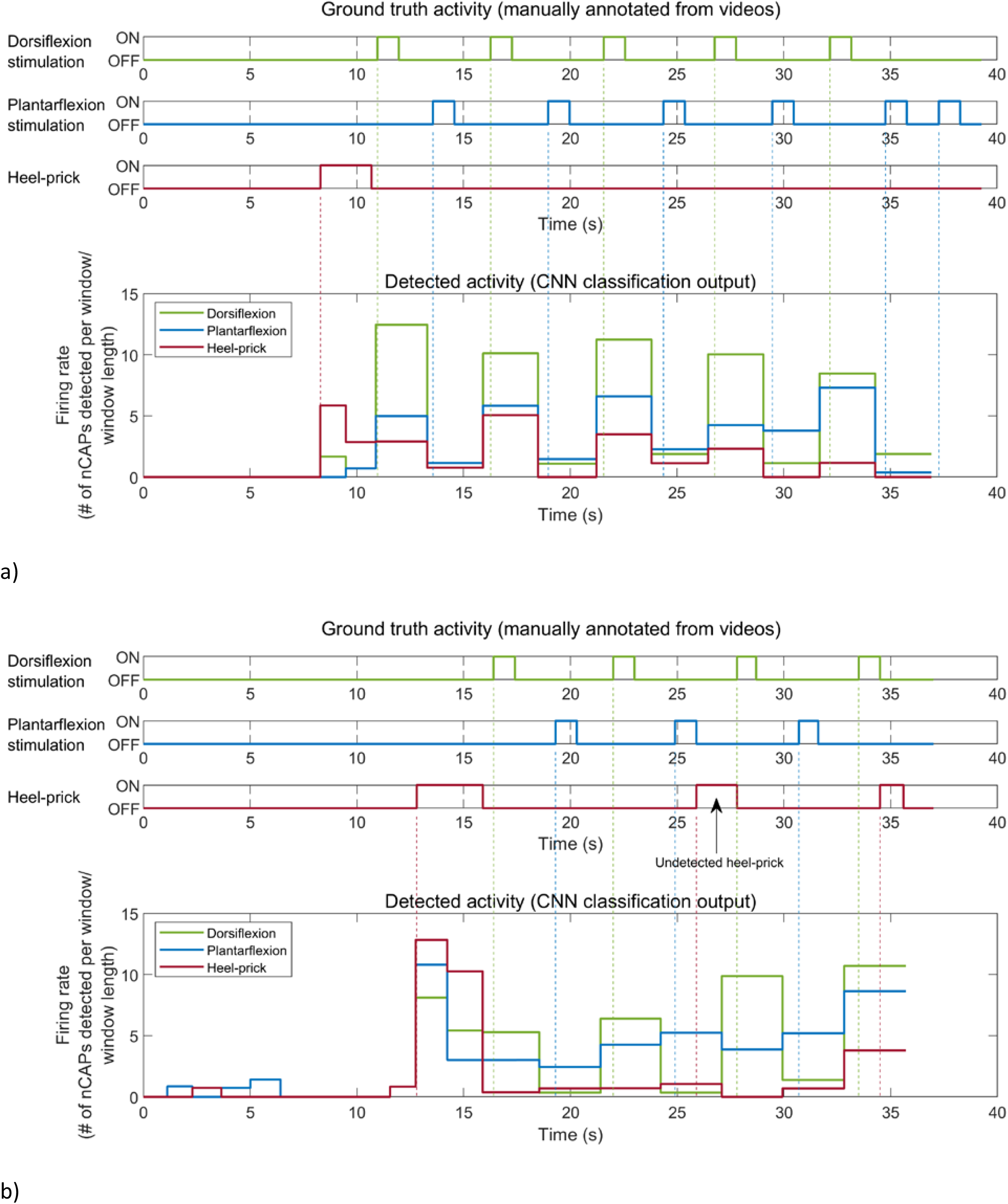
Comparison between ground truth activity and detected activity for 2 example successful trials. Ground truth activity is manually annotated based on video recordings of each trial, and detected activity is based on saved CNN classification outputs. a) Successful trial of FSM A demonstrated in subject A. b) Successful trial of FSM B demonstrated on subject D. Includes one manually annotated heel prick that was not detected by CNN classification.

Table 2 presents the macro F1-score achieved by ESCAPE-NET on the initial datasets acquired off-line from all subjects for which closed-loop FES was attempted. ESCAPE-NET’s off-line performance was evaluated on a test set generated using 3-fold cross validation on each subject’s dataset collected for NN training. Values ranged between 0.47 ± 0.02 and 0.84 ± 0.01. The table additionally reports the macro-F1 scores corresponding to stimulation-evoked trials alone, to give a more direct indication of the expected performance during the closed-loop experiments.

**Table 2:** Macro F1-scores achieved by ESCAPE-NET trained on datasets acquired from subjects for which closed-loop FES was attempted. These results are based on the first training dataset obtained during the experiment. Each network underwent additional training afterwards using updated datasets throughout the experiment.

| Subject | Macro F1-score (offline) | Macro F1-score (stimulated) |
| --- | --- | --- |
| 1 | $0.65 \pm 0.03$ | $0.30 \pm 0.16$ |
| 2 | $0.84 \pm 0.01$ | $0.65 \pm 0.00$ |
| 3 | $0.66 \pm 0.06$ | $0.59 \pm 0.00$ |
| 4 (A) | $0.51 \pm 0.02$ | $0.48 \pm 0.02$ |
| 5 (B) | $0.74 \pm 0.09$ | $0.59 \pm 0.01$ |
| 6 (C) | $0.60 \pm 0.07$ | $0.51 \pm 0.02$ |
| 7 (D) | $0.47 \pm 0.02$ | $0.58 \pm 0.01$ |
| 8 | $0.63 \pm 0.05$ | $0.46 \pm 0.05$ |
| 9 (E) | $0.76 \pm 0.04$ | $0.66 \pm 0.00$ |
| 10 | $0.55 \pm 0.04$ | $0.54 \pm 0.03$ |
| 11 (F) | $0.80 \pm 0.02$ | $0.64 \pm 0.01$ |

### 3.2 Verification of stimulation artefact removal

To verify that neural signal was detected in between stimulation pulses, the average of the resulting spatiotemporal signatures was analyzed and deemed visually similar to the averaged spatiotemporal signatures of manually induced neural activity. Figure 6 illustrates an example of this comparison. In particular, the peak of the nCAP occurring earlier at one longitudinal end of the nerve cuff compared to the other end was an important visual cue that indicated electrical activity travelling along the nerve.

**Figure 6:**
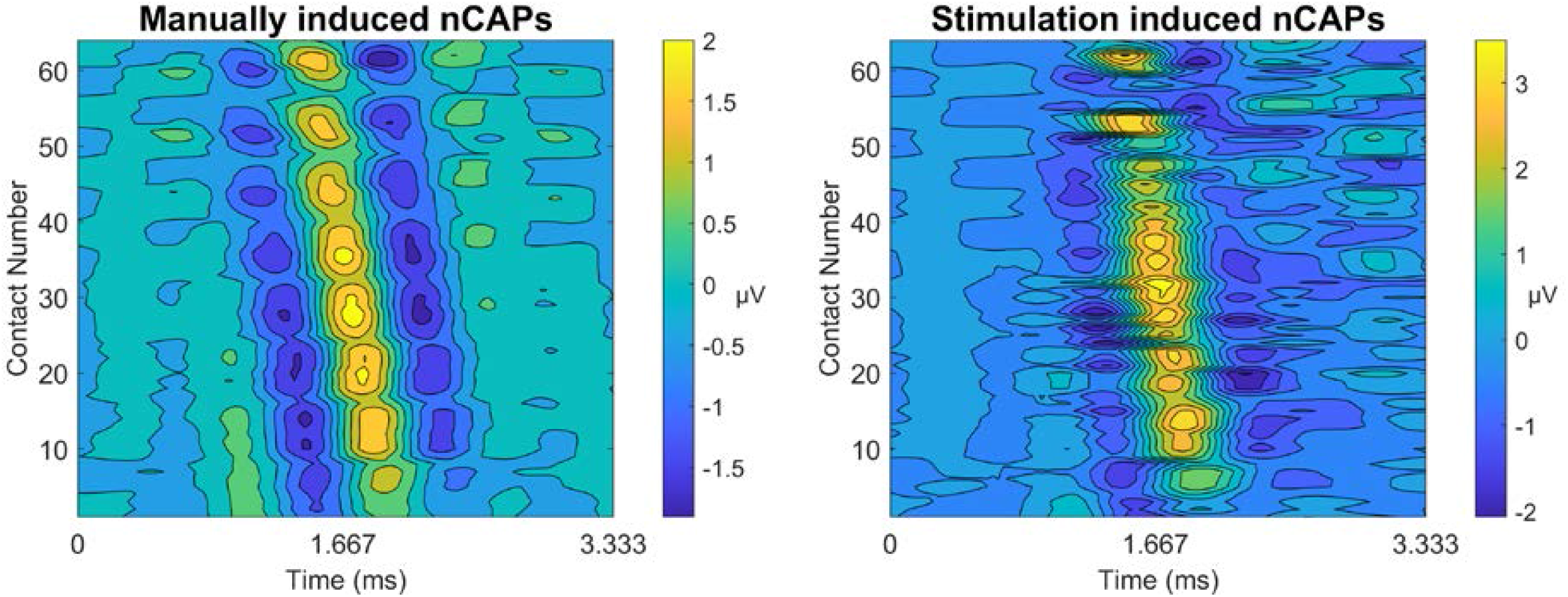
Example of averaged spatiotemporal signatures for nCAPs induced manually (left) and via electrical stimulation (right). The peak region is indicated in yellow, and notably occurs earlier on higher numbered contacts. Numbering was performed such that each ring of contacts is grouped together.

The preprocessing steps taken for nCAP detection was designed to eliminate pulses with high amplitudes outside of the expected range of an nCAP’s amplitude. As nCAPs exhibit amplitudes that are significantly lower than electrically evoked CAPs, it is highly improbable that any electrically evoked CAPs due to stimulation were incorrectly detected as nCAPs. For additional confirmation, the nerve was severed distally, and recording was performed while manually moving the subject’s foot and stimulating the severed nerve as before. No movement was generated by the stimulation, and no nCAPs were detected for the subject used to generate Figure 6 using the same preprocessing steps, hence the absence of a visual depicting spatiotemporal signatures for nCAPs detected after severing the nerve.

## 4 Discussion

This study demonstrated for the first time that classifying individual nCAPs using a CNN can be used to provide feedback information to guide closed-loop electrical stimulation of the nerve. The demonstration was performed in acute *in vivo* experiments on rats using a single custom-designed nerve cuff electrode and commercial off-the-shelf data acquisition and stimulation devices.

### 4.1 Advancements over previous works

Several design choices were made to reduce the invasiveness and increase the number of discriminable neural pathways of our implementation of closed-loop stimulation technology compared to existing approaches.

Previous works demonstrating closed-loop stimulation have used multiple electrodes either to achieve nerve selectivity or to separate recording and stimulation electrodes to diminish stimulation artefacts. Bruns et al. [3] and Holinski et al. [4] both used a microelectrode array for recording; Bruns et al. used 1-3 muscle electrodes per muscle group for stimulation and Holinski et al. used intraspinal wire electrodes for stimulation. This study uses a single nerve cuff electrode capable of simultaneous recording and stimulation.

Using a single nerve cuff electrode yielded complex issues such as signal selectivity, stimulation artefacts, and selective stimulation. This study has achieved recording selectivity and managed stimulation artefacts using signal processing techniques. Minimizing the number of implanted devices in a user reduces the complexity of the required surgical procedure, potentially reducing side effects. Using a single nerve cuff electrode for recording and stimulation also eliminates the need for external sensors (e.g. pressure sensors, accelerometers, position sensors) or muscle stimulators; this is advantageous in the context of reliability, positioning and mounting of external devices [26]. Surgically implanted nerve recording devices have the additional advantage of being externally invisible, a trait that users desire to minimize attention drawn to their condition [27] [28].

Song et al. demonstrated closed-loop FES in rabbits using a single multi-contact nerve cuff electrode on the sciatic nerve for both recording and stimulation [5], discriminating between signals from 2 neural pathways as feedback information. This study discriminates between 3 neural pathways and multiple simultaneously active neural pathways, and classifies individual nCAPs rather than longer time windows.

One of the main advantages of classifying individual nCAPs is the ability to detect multiple simultaneously active neural pathways, which would result in a train of alternating types of nCAPs. Successful trials of FSM B demonstrated the ability to detect activity from two neural pathways happening in temporal proximity. Pricking of the heel would be administered approximately when the foot was fully dorsiflexed or plantarflexed, to induce approximately simultaneous activity in two neural pathways. This work’s ability to differentiate between 3 neural pathways and multiple simultaneously active neural pathways shows its potential applicability to more complex nerves with more neural pathways.

The CNN used in this demonstration is more suitable for implementation on implantable devices compared to the CNN used in [9]. This demonstration used a version of ESCAPE-NET with reduced resource requirements, requiring 7.9x fewer weights and 3.5x fewer floating point operations. Reducing resource requirements reduces computational complexity and therefore the size of required memory devices and power sources required for hardware implementation of the algorithm. Smaller required hardware facilitates implementation on implantable devices. Further reductions of the neural network size are expected to be possible with additional optimization [19].

Apart from the closed-loop demonstration, this study shows that ESCAPE-NET can yield reasonable performance using a different neural interface and contact configuration than previously reported in [10], as reported in Table 2. This finding provides support for the generalizability and reproducibility of the approach. In these experiments, a higher macro F1-score did not always correspond to successful closed-loop FES demonstration. Conversely, a low macro F1-score did not always correspond to failed closed-loop FES demonstration. This may be due to robustness issues, further discussed below, related to the time elapsed between initial dataset collection and closed-loop FES demonstration attempts.

### 4.2 Limitations and avenues for improvement

We encountered robustness issues during this study that should be mitigated or addressed in future studies. These issues resulted in the need to retrain the CNN on newer data, the need to change contacts used for stimulation, unequal amounts of detected nCAPs between the three classes, or occasional failure to achieve closed-loop FES demonstration on particular subjects. Several factors may have contributed to these robustness issues, including nerve damage during the surgical procedure, not removing enough connective tissue from the nerve resulting in poor contact between the electrode and the nerve, inconsistent signal quality and ability to functionally stimulate due to relatively unguided electrode placement, shifting of the electrode position, inconsistent contact coverage around the circumference of the nerve (a region in the middle of the electrode without contacts, and the way the electrode is folded leaves another region without contacts), change in the electrode-tissue interface, insufficient moisture leading to drying of the nerve, influence of anesthesia on signal quality, and electrical interference from nearby devices. Increased stability may be achieved through refining nerve cuff designs to ensure a close and stable interface with the nerve at all contacts and to equalize coverage of contacts across the circumference of the nerve, and using more sophisticated stimulation configurations to better control current flow [29]. Automated methods for evaluating the best contacts (e.g. highest signal-to-noise ratio or contact information metric [9]) and rejecting contacts obtaining poor signal will improve the overall signal quality.

In practice, a user may periodically recalibrate the CNN by providing updated labeled data to either augment or replace the existing dataset and recalibrate appropriate contacts for stimulation. Studies may be performed to optimize the recalibration process so that it can be performed at home by the user in a minimal amount of time with minimal turnaround time for an updated and usable CNN. Sammut et al. simulated natural changes in nerve cuff signals due to connective tissue growth and device rotation, demonstrating that a semi-supervised self learning approach may be used to reduce the frequency of recalibration [30]. To further reduce recalibration frequency, machine learning methods may be explored to autogenerate new data based on predicted trends in signal characteristics to be evaluated using the semi-supervised self learning approach. For example, Shi et al., Saxena et al., and Wang et al. have developed methods of predicting spatiotemporal data using convolutional long short-term memory networks [31], general adversarial networks [32], and recurrent neural networks [33], respectively. To recalibrate stimulation without using the brute-force approach of testing all contacts individually and in combination, methods of reshaping the nerve to move targeted nerve branches closer to the surface [34] or fascicle localisation to facilitate targeted stimulation [35] could be explored.

Our methods for detecting activity in a particular neural pathway was based on either the majority firing rate detected within a time window, or whether a firing rate had exceeded a certain threshold. As a result, the classifier did not need to perform with particularly high accuracy for the FSM to function as expected. Future work should optimize the confidence of a classifier for each window of nerve activity. Manual pricking of the heel using a Von Frey monofilament was difficult to perform in a consistent manner with temporal accuracy, especially while the foot was moving due to nerve stimulation. A structure that can secure the rat limb and apply mechanical pricking of the heel may be able to ensure more consistent manipulation of the subject, similar to what was used by Brunton et al. [36].

A CNN’s high classification ability is often a result of high computational complexity, which translates to larger devices and more computation time due to higher resource requirements. A CNN optimized for implementation on implantable devices should be designed, with a focus on reducing memory and power requirements, while maintaining reasonable classification accuracy for its intended application [19]. This would reduce battery size, hardware form factor and operational latency. Alternatively, a fully implantable neural interfacing device with wireless data communication capabilities could be envisioned, with data processing and stimulation decision-making being performed on another mobile device located outside of the body (or on a cloud platform), and connected in a closed loop with the implanted device [37].

Beyond the efficiency of the CNN itself, the overall system latency in this study was determined by the time needed to read in data from a buffer, process the resulting signal, classify detected nCAPs, and make a corresponding stimulation decision, and was empirically observed to be approximately 3 seconds (as visible in Figure 5). A limitation of the study is that we did not measure the time required for individual operations within this loop. The 3 second buffer duration was appropriate to obtain accurate estimates of firing rates, but the trade-off between accuracy and latency should be further characterized in future work. Another limitation of the study is that we did not consistently document failed attempts. Focus was placed on documenting successful attempts due to the study’s goal of demonstrating a proof of concept.

## 5 Conclusion

Our findings demonstrate that extraneural classification of individual nCAPs is feasible in a closed-loop FES scenario. Individual nCAPs detected in multichannel cuff electrode recordings were classified using the ESCAPE-NET CNN, providing activity levels of the sciatic nerve’s three branches that determined the type of closed-loop stimulation to deliver. This method detects multiple neural pathways firing in temporal proximity, showing potential for applicability to nerves with more neural pathways. Design choices made to minimize invasiveness include the use of a single electrode for simultaneous recording and stimulation and use of a modified version of ESCAPE-NET with reduced resource requirements [9]. These results constitute an important step towards implementing closed-loop FES in clinical applications. Future work should focus on increasing robustness and conducting chronic implantation studies to support translation to humans. Beyond the ability to restore nuanced motor function in paralyzed limbs, closed-loop neuromodulation technology may be used to integrate sensory feedback into prosthetic limbs and to manage ailments such as chronic pain, diabetes, or incontinence.

## 6 Acknowledgments

This work was supported by the Centre for Analytics and Artificial Intelligence Engineering Seed program at the University of Toronto.

## Notes

### Competing Interest Statement

The authors have declared no competing interest.

